# The SPEECHLESS-induced stomatal increase is required for the salt tolerance of oil palm

**DOI:** 10.1101/2021.12.09.471966

**Authors:** Zhuojun Song, Le Wang, Chong Cheong Lai, Zituo Yang, May Lee, Gen Hua Yue

## Abstract

Oil palm is the most productive oil producing plant. Salt stress leads to growth damage and decrease in yield of oil palm. However, the physiological responses of oil palm to salt stress and their underlying mechanisms are not clear. RNA-Seq for leaf samples from young palms challenged under three levels of salts (100, 250 and 500 mM NaCl) and control for 14 days was conducted. Diverse signalling pathways were involved in responses to different levels of salt stress. All the three levels of salt stress activated *EgSPCH* expression and induced stomatal density of oil palm, which was contrasting to that in *Arabidopsis*. Under strong salt stress group, oil palm removed excessive salt via stomata. Overexpression of *EgSPCH* in *Arabidopsis* increased the stomatal production but lowered the salt tolerance. These data suggest that in oil palm, salt activates *EgSPCH* to generate more stomata in response to salt stress. Our results shed a light on the cellular response to salt stress of oil palm and provide new insights into the mechanisms of different salt-induced stomatal development between halophytes and glycophytes.

## INTRODUCTION

Oil palm (*Elaeis guineensis*, Jacq.) produces the highest yields of plant oil (Corley and Tinker, 2008). Due to the negative effects of oil palm expansion, such as deforestation and decreasing biodiversity, sustainable plantation and management is the way to increase oil production and minimize the damage to environment (Fitzherbert et al., 2008). Oil palm is cultivated in tropical areas of Asia, Africa and America (Corley and Tinker, 2008) where many coastal soils of those areas are salinized due to tidal waters (Henry and Wan, 2012).The fresh fruit bunch (FFB) yields of oil palm dramatically decreased on the saline soils (Henry and Wan, 2012).Therefore, the genetic improvement by selecting salt-tolerant oil palm varieties is important for sustainable palm oil production (Corley and Tinker, 2008). However, not much is known about the molecular mechanism underlying salt tolerance in oil palm.

Over the past decade, the molecular mechanisms of salt-tolerance have been largely studied in *Arabidopsis* and agronomic plant species, such as rice (Kumar et al., 2013; Zhang et al., 2021). Salt stress can directly change the biological compounds physically or chemically in plant cells, which cause cellar response (Zhang et al., 2021). Furthermore, salt stress leads to ionic stress, secondary stresses and osmotic stress and oxidative stress, thereby triggering multiple complex signalling pathways (Yang and Guo, 2018). The leucine-rich repeat extensins (LRX)-Raf like kinase (RALF)-FERONIA (FER) module is important for cell wall integrity and cell wall associated biological processes (Feng et al., 2018). In plants, high salinity disrupts the cross-link between pectin and LRXs, and the interaction between LRXs and RALFs, resulting in cell bursting during growth under salt stress (Zhao et al., 2018). Salt stress triggers cytosolic Ca^2+^ signal, which activates the Na^+^ homeostasis required Salt Overly Sensitive (SOS) signalling pathway ultimately, H^+^-ATPase is activated and Na^+^ is exported via Na^+^/H^+^ exchanger driven by H^+^-ATPase (Kumar et al., 2013; Zhang et al., 2021). Many other genes are also important in ionic stress signalling pathway. They repress the salt sensory system, limit the salt absorption and transportation in plants, regulate root and leaf development and adjust the ionic balance of cells to raise up the salt tolerance (Munns, 2005; Deinlein et al., 2014). Transcription factors (TFs) play key roles in the salt stress tolerance of plants. They are differentially expressed during salt stress, which consequently regulate the transcription of various downstream genes that are involved in salt tolerance (Golldack et al., 2011). The most well-known salt tolerance associated TFs, including basic leucine zipper (bZIP), basic helix-loop-helix (bHLH), MYB, WRKY, APETALA2 and NAC (Zhang et al., 2006; Golldack et al., 2011; Van Zelm et al., 2020). Among the TFs, a bHLH transcription factor SPEECHLESS (SPCH) serves as a master regulator of cell development in response to environmental changes (Lau et al., 2014). SPCH binds to ~ 4.5% of genes in *Arabidopsis*, including key genes in abiotic stress and hormonal stress signalling pathway (Lau et al., 2014). The function of SPCH in stomatal initiation is conserved in both dicots and monocots (Lampard et al., 2008; Wu et al., 2019). Under salt stress, the expression of SPCH was repressed by upstream transcriptional factors and mitogen-activated protein kinase (MAPK) signalling pathway, resulting in the reduction of stomatal production in order to avoid water loss (Kumari et al., 2014). Although these studies provide novel knowledges and new insights of the regulatory networks of salt tolerance, the complexity of salt resistance, the genetical divergence of different species and the diversity of environments make it difficult to understand the particular mechanisms of other plants in response to salt stress (Van Zelm et al., 2020).

Only very few studies show the physiological and proteomic changes of palms in response to salt stress. In oil palm seedlings subjected to salt stress, the content of Na^+^ and proline increased, and the cell membrane was injured in samples treated by the highest salinity at 200 mM NaCl. On the contrary, photosynthetic and growth rate were reduced (Cha-Um et al., 2010). A proteome study of date palm suggests that ATP synthase and RubisCO activase are significantly changed during salt stress (El Rabey et al., 2016), indicating the importance of biosynthesis for salt tolerance. These studies show the physiological responses of palms under salt stress. However, the cellular level response and the molecular mechanisms of the salt tolerance of palms are still unknown.

The purpose of this study was to investigate the salt response of oil palm on cellular level and identify the critical regulators and signalling pathways involved in salt-tolerance. Herein, we found that oil palm exhibits diverse biological strategy in response to different level of salt stress. Furthermore, we found salt stress induced converse regulation of *SPEECHLESS* expression in oil palm and *Arabidopsis*, which leads to the reverse stomatal response. EgSPEECHLESS putatively regulates the expression of 41% of the DEGs. Our study shed a light on the molecular mechanism that explain the different physiological and cellular responses to salt between tree crops and herb crops.

## RESULTS

### Morphological and physiological responses to salt tolerance

Oil palm seedlings with same developmental stage were selected for salt stress assay with daily watering 150 mL of the following four gradient NaCl concentrations: 0 mM (Mock, water only) 100 mM, 250 mM and 500 mM for 14 days. Rescue-assay with watering was performed for another 14 days. Common plant stress responses, including leaf tip necrosis, leaf yellowing and wilting, were observed in all the salt treated samples (Figure 1A). In addition, the roots of salt treated oil palms shrank or even rotted after 14 days (Figure 1A). With the increasing of salt concentration, the above responses of leaf and roots were enhanced (Figure 1A, C). Interestingly, salt emitted and crystalized on leaf epidermal of oil palms treated with high concentration of salt at 250 mM and 500 mM (Figure 1C, Supplemental Figure S1), suggesting that under strong salt stress, oil palm discharges the absorbed salt by transpiration stream via stomata. This physiological reaction was found in halophytes (Robinson et al., 1997) but was rare in non-halophytes, indicating that oil palm may has high salt tolerance as a non-halophyte.

**Figure 1.**
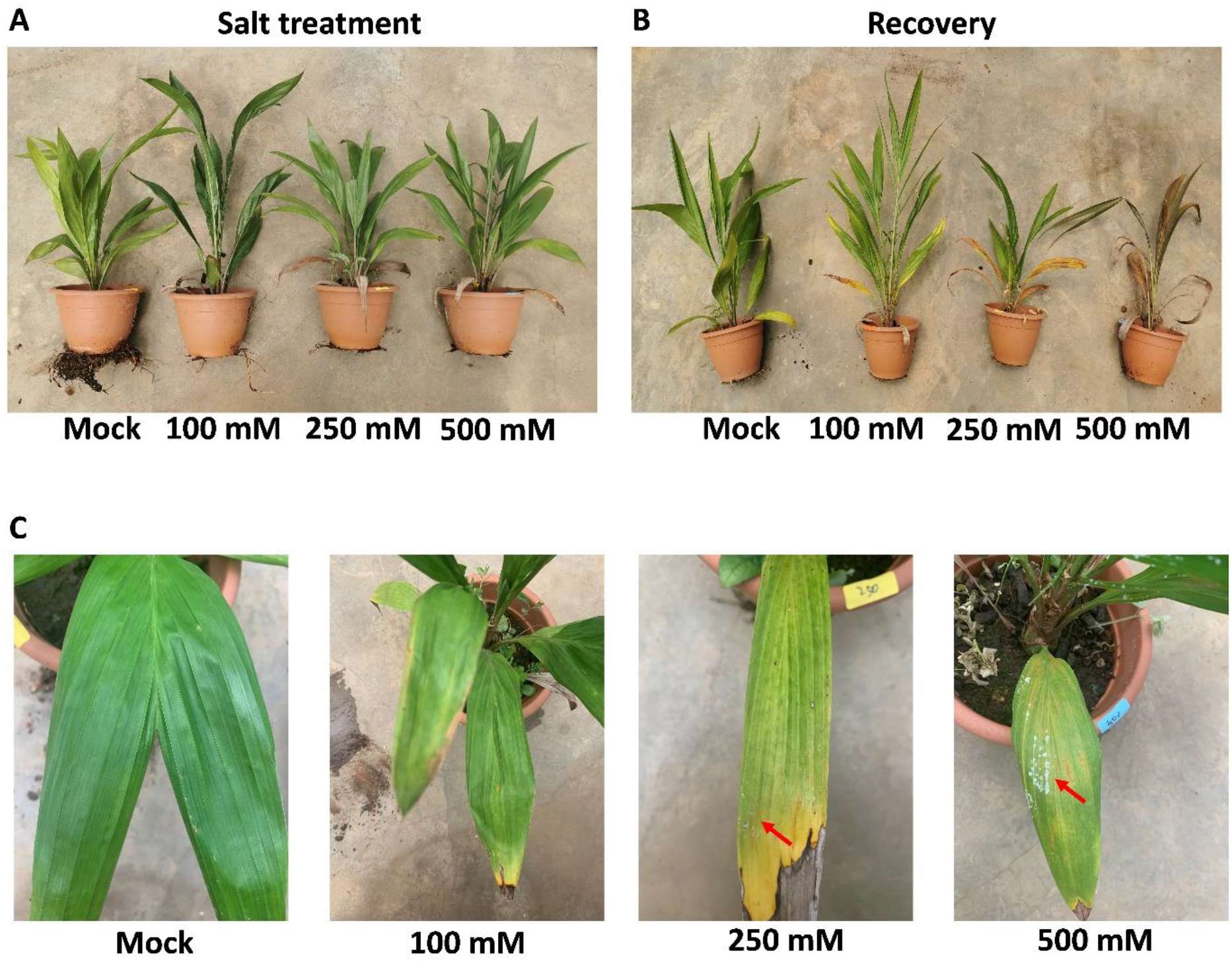
Phenotypical changes of oil palm seedlings in response to different level of salt stress. A, ~2-year-old oil palm seedlings were treated daily with either 150 mL water (Mock) or 150 mL NaCl with four gradient concentration: 100 mM, 250 mM and 500 mM for 14 days. Four seedlings were used in each group as biological repeats. B, all the seedlings from (A) were recovered with 150 mL water daily for another 14 days. C, The leaves of oil palms from (A). Red arrows indicate the salt crystals emitted from leaf surface.

To investigate the effect of salt stress on later growth of plants, rescue assay was performed by giving all the samples 150 mL water daily for another 14 days. Plants treated with 2-weeks of salt with 100 mM NaCl and 250 mM NaCl survived after rescue assay. However, 500 mM NaCl was lethal to long term growth of oil palm (Figure 1B). These results indicate that oil palms show diverse physiological responses to different level of salt stress.

### DEGs of oil palm in response to different level of salt stress

Average cleaned reads of 46.2, 35.0, 59.4 and 35.9 million were obtained and from the Mock, 100 mM, 250 mM and 500 mM NaCl groups, respectively (Supplementary Table S7). A total of 363, 242 and 433 DEGs were identified from salt stress groups (100, 250 and 500 mM NaCl, Figure 2). In detail, 86 down-regulated and 277 up-regulated DEGs were identified in 100 mM NaCl group (Figure 2A), 155 down-regulated and 87 up-regulated DEGs were identified in 250 mM NaCl group (Figure 2A), 249 down-regulated and 184 up-regulated DEGs were identified in 500 mM NaCl group (Figure 2A). PCA and hierarchical clustering analyses were performed. The control and salt treatment groups were clearly differentiated and showed substantial differences (Figure 3, Supplemental Figure S2). In addition, three DEGs (*EgSPCH*, *EgPAT1* and *EgRPS3*) were up-regulated in all the salt treatment groups (Figure 2A). EgSPCH is a homolog of Arabidopsis SPEECHLESS, which is a bHLH transcription regulator that directly controls stomatal development and regulates the expression of thousands of genes (Lau et al., 2014). Both PAT1 and RPS3 are expressed in chloroplast. *PAT1* decays ABA responsive genes thereby regulating salt tolerance (Zuo et al., 2021). *EgRPS3* is required for plant pathogen resistance (Bisgrove et al., 1994). These data suggest that although only a few of DEGs were overlapped across different level of salt stress, light-induced biological process and stomatal development are required in general defence of oil palm in response to different level of salt stress.

**Figure 2.**
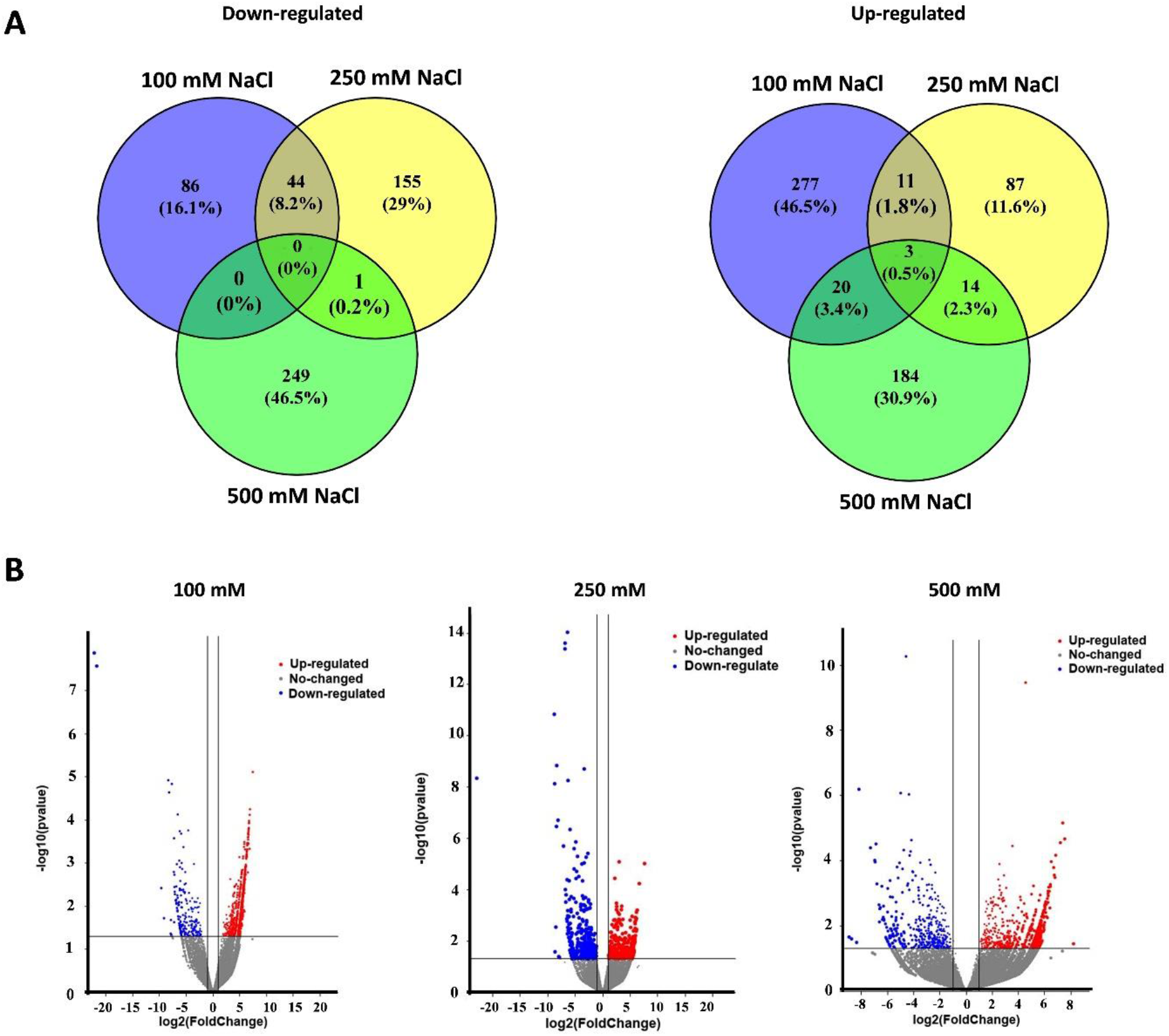
Comparison of differentially expressed genes (DEGs) in the young leaves of oil palm seedlings under three level of salt stress: 100 mM NaCl, 250 mM NaCl and 500 mM NaCl. A, Up- and down-regulated DEGs showed by Venn diagrams. B, *P*-value and log2foldchange (Log2FC) of DEGs under 100 mM, 250 mM and 500 mM NaCl showed by volcano plots.

**Figure 3.**
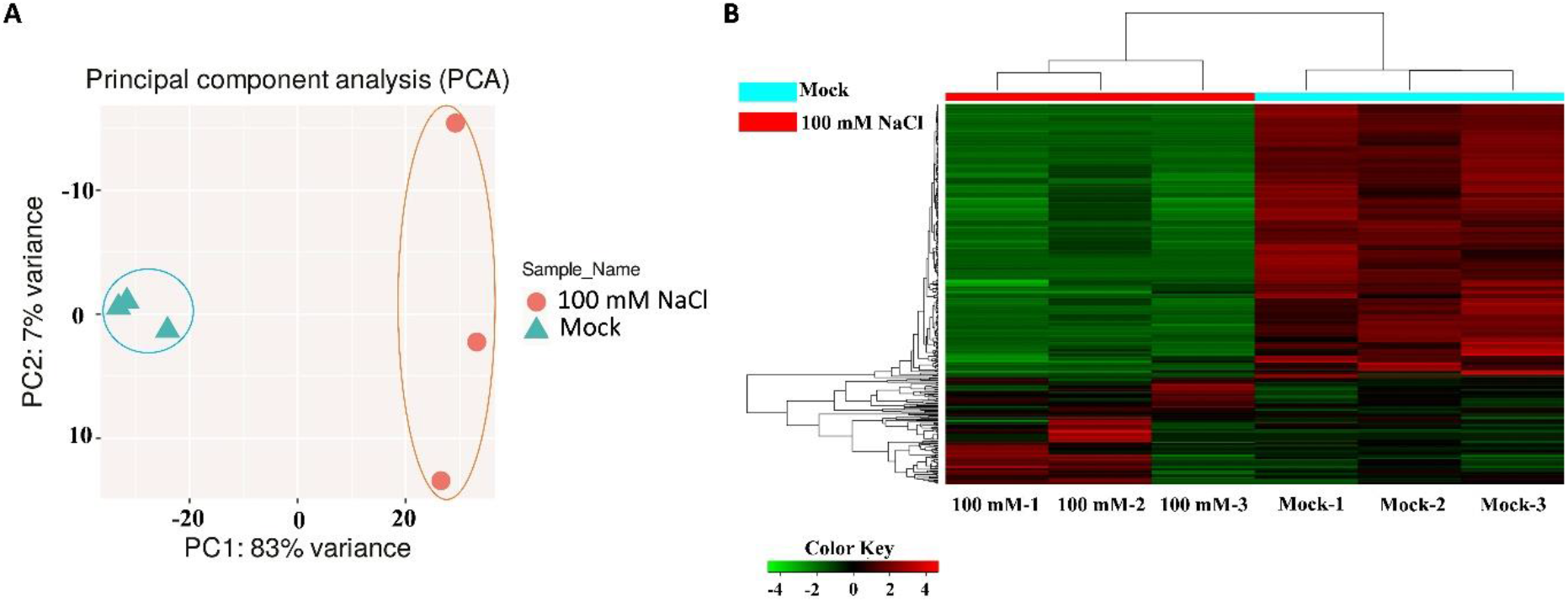
The diversities of differentially expressed genes (DEGs) in response to 100 mM NaCl. A, Principal component analysis among samples of the 100 mM NaCl and Mock groups based on randomly selected DEGs. B, Hierarchical clustering among samples of the 100 mM NaCl and Mock groups based on randomly selected DEGs. 100 mM1, X100 mM2 and X100 mM3 are three biological repeats of 100 mM NaCl salt stress group, while Mock1, Mock2 and Mock3 are three biological repeats of Mock group. Up-regulated DEGs and Down-regulated DEGs are represented by red and green bars, respectively.

### Oil palms exhibit diverse biological strategies in response to different levels of salt stress

Analysis of GO enrichment showed that in samples treated by low level of salt (100 mM NaCl), defense/stress response, metabolic process and plant development were the main signaling pathways of the DEGs (Table 1, Figure 4A). Most of the salt response related DEGs were up-regulated while most of the DEGs in terms of response to biotic stimulus (bacterium and fungi) and other abiotic responses were down-regulated (Table 1, Figure 4A). In addition, genes regulating other development such as seed, ovule and roots, were down-regulated (Table 1), implying the metabolic and cell developmental compensation in response to salt stress by sacrificing other defense systems and development events.

**Table 1.** Selected DEGs and their KEGG pathways in response to 100 mM NaCl challenge in the young rosette leaves of oil palm seedlings

| Gene | Annotation | Subsignaling pathways | Expression |
| --- | --- | --- | --- |
| Defense response and stress response (35.9%) |  |  |  |
| LOC105040031 | acts upstream of or within defense response to bacterium | response to stress | Down |
| LOC105036933 | involved in cellular response to salt stress | response to abiotic stimulus | Up |
| LOC105059761 | acts upstream of or within response to salt stress | response to stress | Up |
| LOC105048273 | acts upstream of or within response to salt stress | response to abiotic stimulus | Up |
| LOC105058610 |  | response to stress | Down |
|  | acts upstream of or within cellular response to phosphate starvation |  |  |
| LOC105054129 | acts upstream of or within response to gibberellin | response to endogenous stimulus | Up |
| LOC105050654 | involved in regulation of response to reactive oxygen species | response to stress | Down |
| LOC105040397 | acts upstream of or within response to abiotic stimulus | response to abiotic stimulus | Down |
| catabolic and other metabolic process (34.3%) |  |  |  |
| LOC105060222 | acts upstream of or within proteasomal protein catabolic process | catabolic process | Down |
| LOC105060233 | acts upstream of or within ubiquitin-dependent protein catabolic process | other metabolic processes | Down |
| LOC105039973 | enables RNA helicase activity | catalytic activity | Down |
| LOC105054501 | enables ammonia-lyase activity | catalytic activity | Up |
| LOC105041996 | enables serine-type endopeptidase activity | catalytic activity | Down |
| LOC105058751 | enables ubiquitin-protein transferase activity | catalytic activity | Down |
| LOC105041996 | enables serine-type endopeptidase activity | catalytic activity | Up |
| LOC105034218 | enables lysine-tRNA ligase activity | catalytic activity | Down |
| cell and tissue development (29.8%) |  |  |  |
| LOC105040725 | involved in regulation of stomatal complex development | anatomical structure development | Up |
| LOC105043171 | acts upstream of or within regulation of stomatal opening | other cellular processes | Up |
| LOC105038743 | acts upstream of or within embryo development ending in seed dormancy | post-embryonic development | Down |
| LOC105040468 | involved in regulation of photoperiodism, flowering | multicellular organism development | Down |
| LOC105052855 | acts upstream of or within plant ovule development | anatomical structure development | Down |
| LOC105032026 | acts upstream of or within lateral root development | anatomical structure development | Down |
| LOC105055813 | acts upstream of or within root hair elongation | multicellular organism development | Down |
| LOC105058130 | acts upstream of or within xylem and phloem pattern formation | multicellular organism development | Down |

**Figure 4.**
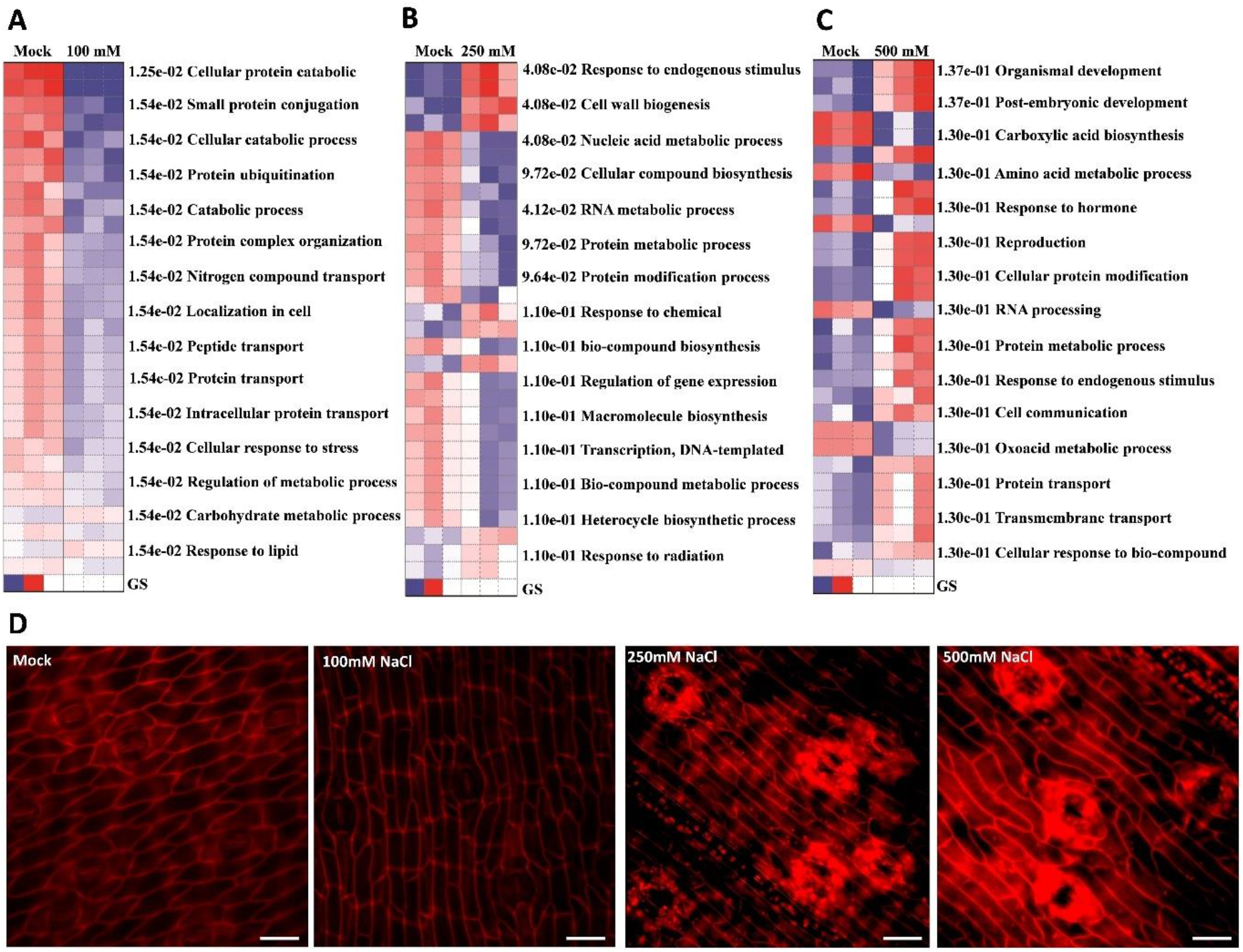
Oil palm exhibits diverse biological responses to different level of salt stress. A-C, Enrichment of gene ontology (GO) of DEGs against salt challenge at the significance level of 0.05 in the young leaves of oil palm seedings under 100 mM NaCl (A), 250 mM NaCl (B) and 500 mM NaCl (C). Up-regulated DEGs and Down-regulated DEGs are represented by red and blue bars. D, Epidermal cells of young leaves sampled from Mock(d), 100 mM NaCl(e), 250 mM NaCl(f) and 500 mM NaCl(g) staining by PI were showed, scale bar = 25 μm.

In 250 mM NaCl group, biosynthesis and metabolic process contribute equally (48.4% and 44.4%) in response to salt stress (Supplemental Table S2, Figure 4B) where secondary metabolites synthesis and cell wall biogenesis were dominant in the regulatory signaling pathways (Supplemental Table S2, Figure 4B). The accumulation of flavonoid and chalcone during salt stress were largely found in other crops as they are important for plant salt tolerance by maintaining reactive oxygen species (ROS, (Lijuan et al., 2015; Chen et al., 2019). Positive regulators of flavonoid (LOC105055971, LOC105054663) and chalcone (LOC105050962) synthesis were up-regulated (Supplemental Table S2). The cell wall is a crucial component of the plant cell which is highly dynamic and quickly responsive to abiotic stimulus. The maintenance of cell wall homeostasis is essential for the stress tolerance of plant cells (Zagorchev et al., 2014; Zhao et al., 2018). The epidermal cells of fresh young leaves sampled from mock and salt stress groups showed that under salt stress, the epidermises consisted of more and longer pavement cells (Figure 4D). The cell wall integrity of 100 mM NaCl group was comparable with the Mock group (Figure 4D), while in the 250 mM NaCl and 500 mM NaCl group, the epidermal cells, especially the guard cells and their surrounding pavement cells were largely damaged and propidium iodide (PI) permeated into the cytosol of these necrotic cells (Figure 4D). Unlike the epidermal cells in Mock and 100 mM NaCl group, which were linearly distributed, the epidermal cells in 250 mM and 500 mM were tortile (Figure 4D). These data suggest that high salinity soil is harmful to the cell integrity. Thus, in response to the high salinity, the oil palm increased the cell wall biogenesis in order to maintain homeostasis. (Supplemental Table S2, Figure 4B).

In samples that have undergone high salt stress (500 mM NaCl), DNA & RNA processing and amino acids & sugar metabolisms are key pathways in response to salt stress. The expression of some genes associated with starch and glycogen synthesis (LOC105047182, LOC105058934 etc.) are inhibited (Supplemental Figure S3, Figure 4C), which might lead to the reduction of starch accumulation during salt stress. This result is in agreement with the previous study in rice (Chen et al., 2007). Most of DNA damage repair genes were up-regulated, suggesting that high salinity may cause severe DNA damage and thus activating the DNA repair system of oil palms (Supplemental Table S3, Figure 4C).

Interestingly, genes that regulate stomatal development and stomatal movement were up-regulated in all the salt treatment groups (Table 1, Supplemental Figure S2–3), implying the importance of stomata in salt tolerance. Taken together, our data suggest oil palm activates its salt tolerance signaling pathways in response to low salt stress. Secondary metabolism synthesis and cell wall biogenesis were enhanced to improve the cell integrity of samples treated by 250 mM NaCl. There was possible DNA damage in high salinity samples (500 mM NaCl) and DNA damage repair pathway was significantly activated. Importantly, stomatal development and stomatal movement were required for salt tolerance of oil palm in response to both low and high salt stress.

### The balance of stomatal development and movement are required for salt tolerance

Stomata is an ion-sensitive valve that control gas exchange and water emission thereby playing essential roles in abiotic stress tolerance (Vahisalu et al., 2008). Chloride channel (CLC) family functions in salt tolerance by regulating stomatal movement via controlling nitrate homeostasis and pH adjustment in *Arabidopsis* (Jossier et al., 2010). In rice, DST (DROUGHT AND SALT TOLERANCE) regulates salt tolerance by controlling stomatal movement via modulating H2O2 homeostasis (Huang et al., 2009). In our study, DEGs in terms of stomatal development and movement were identified in all the salt stress groups (Table 1, Supplemental Table S2–3), suggesting the importance of stomata in salt resistance. To understand how the stomata contributes to the salt tolerance of oil palm, the stomatal density and stomatal aperture of samples from Mock and salt stress groups were monitored (Figure 5). In salt-treated groups, the stomatal apertures were significantly smaller than that in mock group (*p* < 0.01). Furthermore, in higher salinity groups (250 mM and 500 mM NaCl), the stomatal aperture is smaller than that in low salinity group (100 mM NaCl). In salt treatment groups, the stomatal density was higher than that in control group, but there was no difference between these salt stress groups (Figure 5). These data suggest that salt-induced osmotic stress strongly represses the stomatal opening but activates stomatal development (Figure 5). Our data supports the previous studies in rice and *Arabidopsis* that plants reduce the stomatal apertures to limit water loss and reduce transpiration under salt stress (Huang et al., 2009; Jossier et al., 2010). However, our findings that salt stress induces higher stomatal density differs with previous studies in other herbaceous crops, which found that lower stomatal density facilitates the salinity adaption (Huang et al., 2009; Orsini et al., 2012). Interestingly, salt also induces the reduced stomatal aperture and the increase of stomatal density in a ligneous plant *Populus alba L*, where the salt tolerant line 14P11 shows higher stomatal density and smaller stomatal size (Abbruzzese et al., 2009). According to our data, oil palm exhibited halophyte-like salt emission and could survive under 100 mM NaCl for long periods (Figure 1). Higher stomatal density may facilitate the emission of salt along with transpiration, whilst at the same time, the smaller stomatal aperture is beneficial in restricting water loss. Our data suggest that the balance between stomatal density and stomatal movement is required for salt tolerance of oil palm.

**Figure 5.**
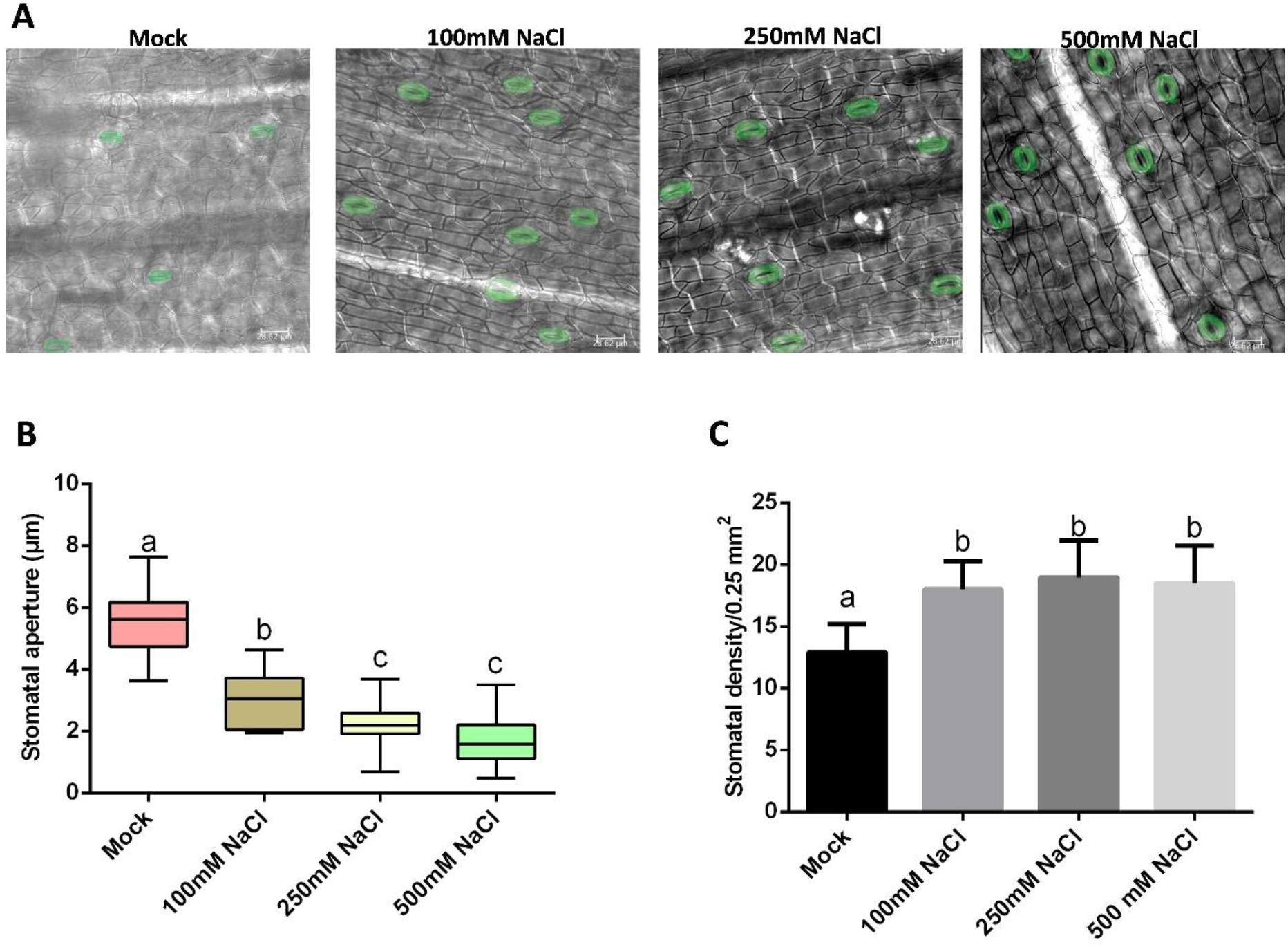
Salt stress represses stomatal opening and activates stomatal production in oil palm. A, Stomata of oil palm in Mock, 100 mM NaCl, 250 mM NaCl and 500 mM NaCl groups, stomata are green coloured, scale bar = 30 μm. B, Stomatal aperture of samples from (A). C, Stomatal density of samples from (A). Values are mean ± SD; n = 20. One-way ANOVA with post hoc Tukey HSD; *p* < 0.01. Samples were treated daily with 150 mL of either water (Mock) or NaCl for 14 days.

### The overexpression of oil palm SPEECHLESS facilitates stomatal development and decrease salt tolerance in *Arabidopsis*

SPEEECHLESS is a key bHLH transcription factor that binds and regulate thousands of genes, and is also a master regulator in stomata initiation (Lau et al., 2014). However, the association between SPEECHLESS and salt tolerance is unclear. In our study, the expression of oil palm *SPEECHLESS* (*EgSPCH*/LOC105040725), which is a homolog gene of *AtSPEECHLESS*, was up-regulated in all the salt treatment groups (Table 1-3, Supplemental Figure S3). In order to determine the function of EgSPCH in stomatal development and salt tolerance, the CDS of *EgSPCH* was cloned into pBGW541 vector driven by the 35S promoter. The plasmid was then transformed into *Arabidopsis* and the transformation was validated by microscopy and PCR (Figure 6C). Like AtSPEECHLESS (AtSPCH), EgSPCH was also localized in the nucleus of epidermal cells (Figure 6A). The introduction of *35S:EgSPCH* significantly increased the stomatal production in *Arabidopsis* (Figure 7C), while both Col-0 and *35S:EgSPCH* exhibited decreased stomata in 150 mM NaCl treatment (Figure 7C, D). The result of a salinity assay showed that the 35S:EgSPCH-YFP plants had a lower salt tolerance (Figure 7A, B). These results indicate the similarity of EgSPCH and AtSPCH in facilitating stomatal development. Our results in 35S:EgSPCH-YFP plants is in agreement with a previous study where the high salinity stress inhibits the growth and stomatal development of *Arabidopsis* (Kumari et al., 2014). However, in oil palm, *SPCH* showed an opposite transcriptional response, where it was activated by salt. This was likely induced by unknown upstream signaling pathway that activates *SPCH* expression in oil palm.

**Figure 6.**
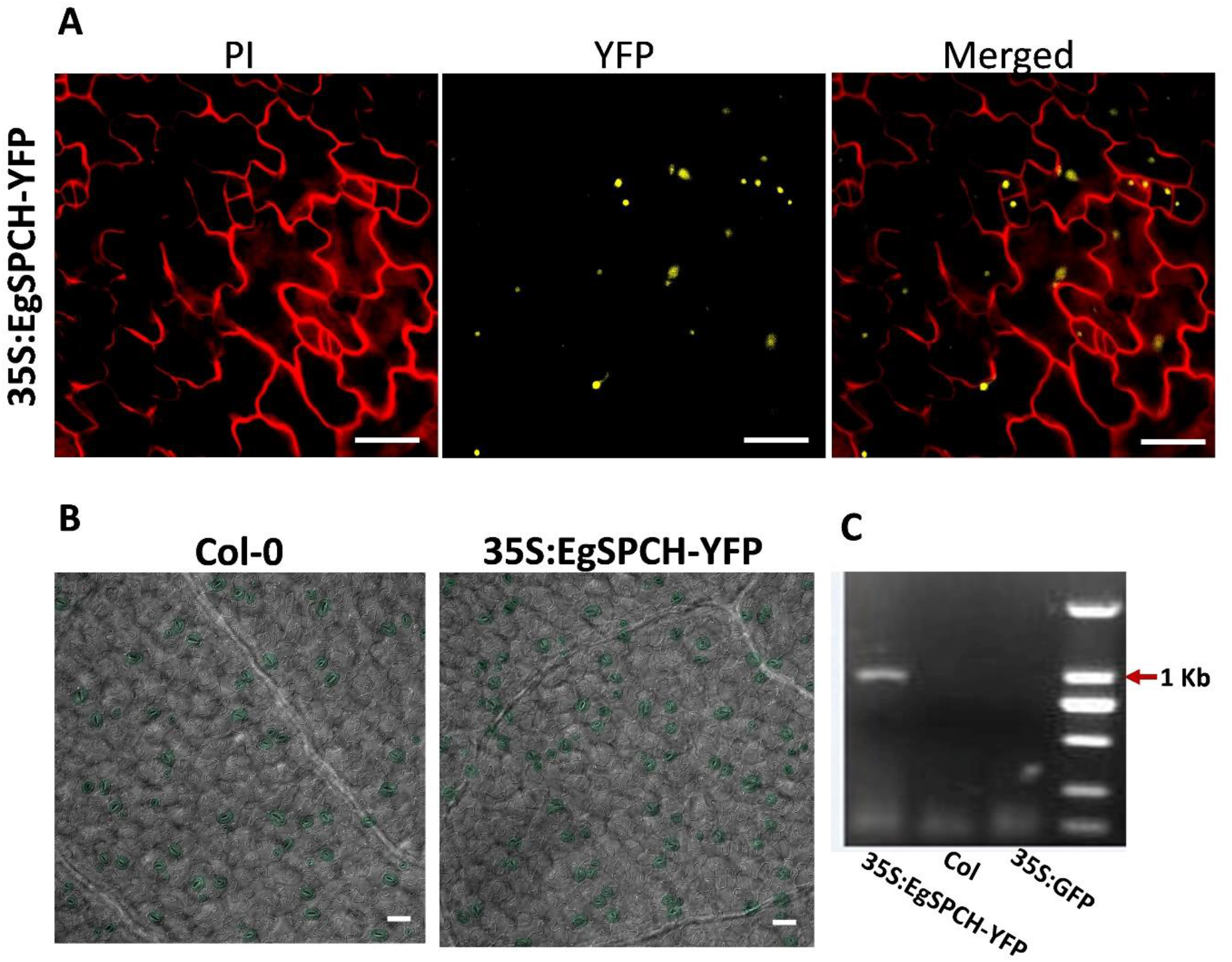
Subcellular localization of EgSPCH and the phenotype of transgenic *35S:EgSPCH-YFP* in *Arabidopsis*. A, Subcellular localization of *35S-EgSPCH-YFP* in 3dpg abaxial cotyledons. PI staining was used for cell outline. B, Stomata of Col-0 and *35S-EgSPCH-YFP* in 3dpg abaxial cotyledons, scale bar = 20 μm. C, Validation of transgenic *35S-EgSPCH-YFP* using PCR genotyping, a product including 3’EgSPCH and 5’YFP with 1022 bp was amplified.

**Figure 7.**
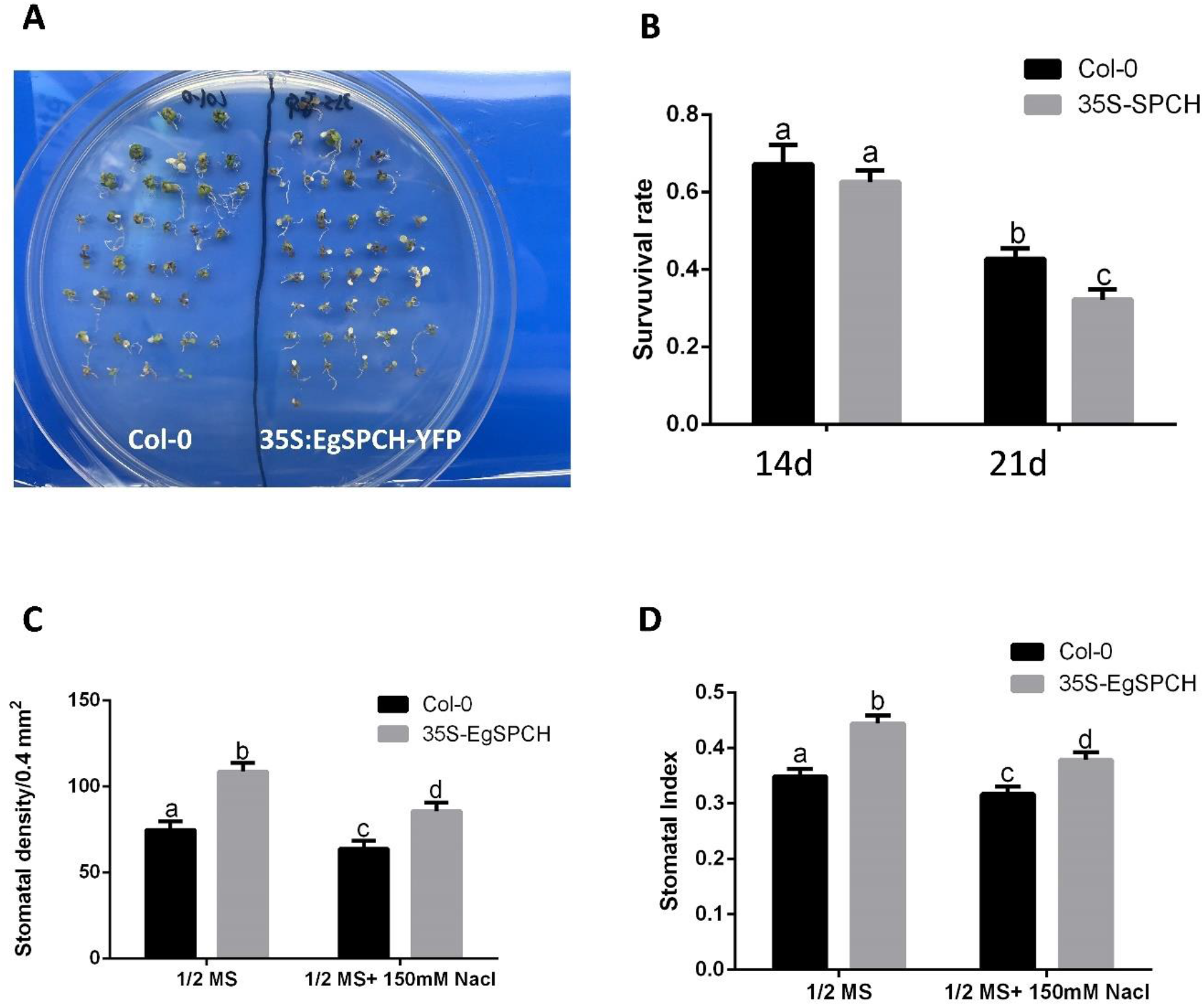
Overexpression of EgSPCH increase stomatal production and decrease salt tolerance in *Arabidopsis*. A, Col-0 and *35S-EgSPCH-YFP* seeds were germinated and grown on ½ MS plates for 7days, after which, they were transferred to ½ MS+ 150 mM NaCl plates for 14 more days (21 dpg). B, Survival rate of Col-0 and 35S-EgSPCH-YFP seedlings at 14 dpg and 21 dpg. n = 40. C, Stomatal density of samples from (A) at 14 dpg. D, Stomatal index of samples from (A) at 14 dpg. Values are mean ± SD; n = 20. One-way ANOVA with post hoc Tukey HSD; p < 0.01

### SPCH is a key molecular switch of transcriptomic response to salt stress

In *Arabidopsis*, SPCH directly controls the transcription of thousands of genes, including key regulators in abiotic stress responsiveness, hormonal signaling and developmental processes (Lau et al., 2014). To identify the effect of alternative EgSPCH expression during salt stress, our DEGs were compared with the chromatin immunoprecipitation (ChIP) sequencing dataset of AtSPCH targets in *Arabidopsis* (Lau et al., 2014). In total, 40.9% of DEGs (with 60.1% and 38.6% of up- and down-regulated genes, respectively) were putative targets of EgSPCH (Figure 8A, Supplemental Table S6). Gene Ontology (GO) terms for genes involved in salt-tolerance, including hormonal and abiotic stress stimulus, developmental processes, organic compound biogenesis and metabolic processes were significantly enriched (Figure 8C). In addition, SPCH plays a key role in transcriptional regulatory cascade of the salt tolerance of plants via controlling the expression of other transcription factors (Lau et al., 2014). In this study, EgSPCH putatively binds to bHLH, MYB, C2H2, NAC, bZIP and many other transcription factors (Figure 8B, Supplemental Table S4). The high percentage of EgSPCH targets among DEGs suggests that EgSPCH is a key transcriptional switch for the salt tolerance of oil palm, EgSPCH and its targets were highly responsive to salt stress, thereby regulating multiple downstream signaling pathways.

**Figure 8.**
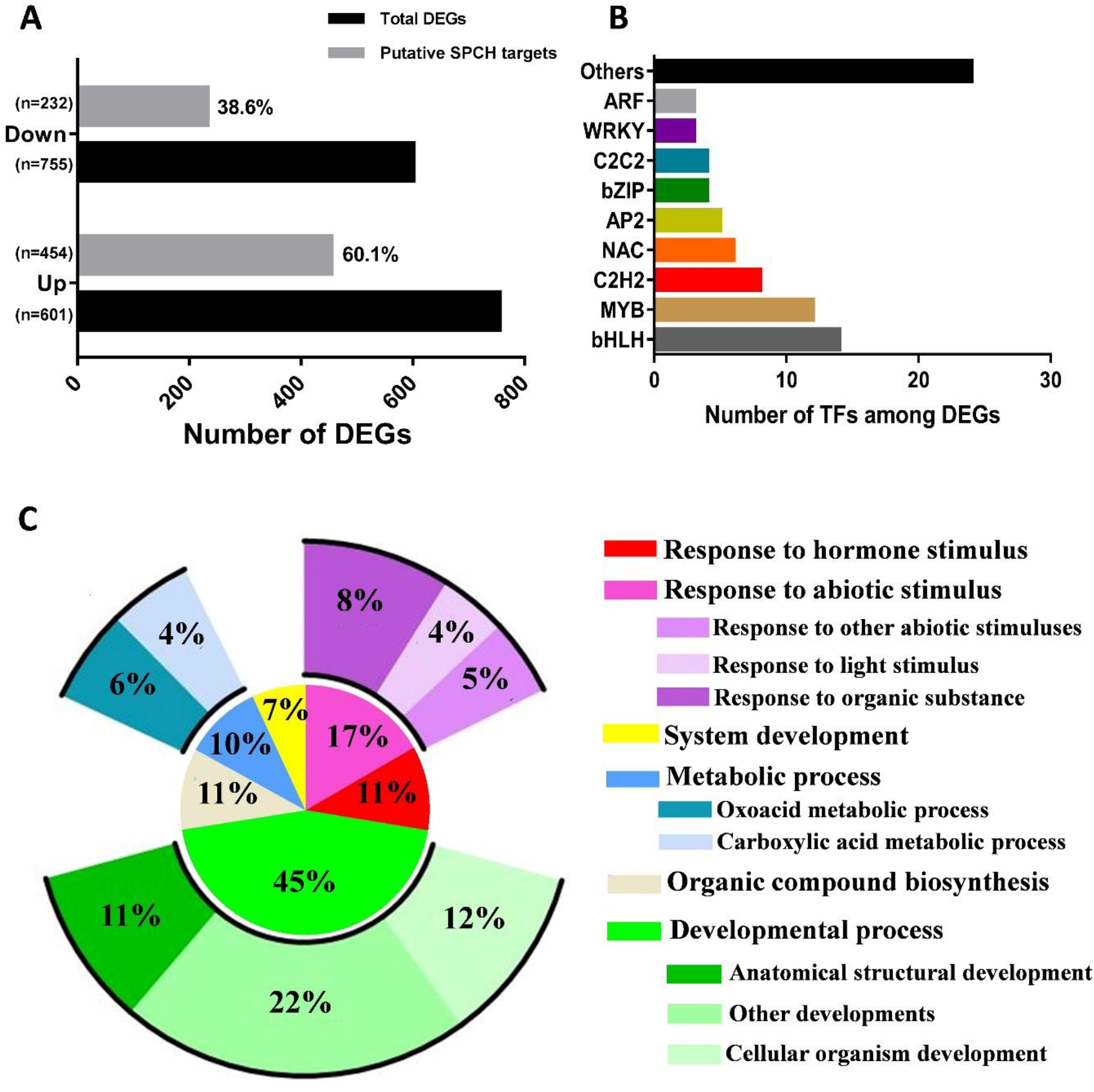
The EgSPCH targets among DEGs are involved in multiple salt tolerance biological processes. A, Percentage of putative EgSPCH targets DEGs in RNA-seq analysis. B, Enriched GO terms of EgSPCH targets

## DISCUSSION

### Similar salt stress response of stomata between oil palm and halophytes

Stomata are minute openings found in the epidermis of the plants, which control CO_2_ intake for photosynthesis and regulates water loss. Stomata consist of pairs of guard cells, which are required for stomatal movement (Hetherington and Woodward, 2003). The ATP driven proton pumps in guard cells are key elements for stomatal movement, which are highly sensitive to various environmental changes (Hetherington and Woodward, 2003). The dynamic changes of stomatal development and movement in halophytes under salt stress have received attention. In non-halophytes, salt stress causes increase in ABA biosynthesis, H_2_O_2_ accumulation and K+ availability reduction, which represses stomatal development and induces stomatal closure (Hedrich and Shabala, 2018). However, the stomata of naturally salt tolerant halophytes function well in high salinity that would kill most other plants (Hedrich and Shabala, 2018). ABA content remain constant in the leaves of halophyte, on the other hand, polyphenols, specifically flavanols, accumulate much faster and maintained a higher content level in guard cells of halophytes than in the glycophytes, which are required for guard cell sensitivity to ROS (Watkins et al., 2017). Interestingly, in halophytes, stomata are also pipe for salt discharge (Chen et al., 2019). In our study, although the stomatal aperture was still affected by salt stress (Figure 7C), the stomatal production was not repressed, allowing the salt discharge via stomata (Figure 1C). The salt stress assay showed that oil palms were able to grow well in 100 mM NaCl with no obvious morphological changes and could survive in long periods of 250 mM NaCl treatment, suggesting a relatively higher salt tolerance than glycophytes where 250 mM is lethal (Stepien and Johnson, 2009). Collectively, oil palm exhibited intermediate salt tolerance and physiological response to salt between halophytes and glycophytes, thereby providing the possibility of oil palm transplantation in coastal saline soils via genetic selection.

### DEGs of three levels of salt stress were involved in different signaling pathways

In a previous study, the growth of oil palms was inhibited when exposed to 200 mM NaCl (Cha-Um et al., 2010). However, the repression of growth was not obvious when expose to low levels (50 and 100 mM NaCl) of salinity (Cha-Um et al., 2010). To investigate the cellular responses to a larger salinity gradient and the molecular mechanisms behind them, oil palm seedlings were exposed to three levels of salinities, 100, 250 and 500 mM of NaCl. In general, genes among auxin/ABA induced signaling pathways involved in stress response, plant development and flavonoid biosynthesis were regulated (Supplemental Table S4). At cellular level, pathways involved in stomatal complex development and stomatal movement were significantly regulated (Supplemental Table S4). Although the significant transcriptomic changes were found in all the salt stress groups (Figure 3, Supplemental Figure S2), the DEGs were different and were involved in different signaling pathways (Figure 1, Figure 4, Table 1, Supplemental Figure 2–3). In 100 mM NaCl group, although oil palm seedlings did not exhibit obvious growth repression within 14 days (Figure 1), the transcriptome was largely changed. In 250 mM and 500 mM groups, cell membrane and cell wall were damaged (Figure 4, Supplemental Table S2–3), which was similar to the previous study in oil palm exposed to 200 mM NaCl (Cha-Um et al., 2010). Therefore, the cell wall biosynthesis signalling pathway was activated in higher salinity levels (Supplemental Table S2, S4). However, the exposure to low salinity for a short period is sufficient to activate the cell wall biosynthesis in *Arabidopsis* (Shen et al., 2014), supporting our conclusion that oil palm showed a relatively higher salt tolerance. DNA damage and protein degradation were pronounced under strong salt stress (Ma et al., 2006; Zvanarou et al., 2020). In 500 mM group, many genes involved in DNA repair and protein metabolic signalling pathway were regulated (Supplemental Table S3-4). The transcriptional analysis of oil palm rosette leaves under different salinity levels suggests that oil palm use diverse biological strategies in response to salt stress. Among those strategies, stomatal development and movement contribute to the cellular response to multiple levels of salinity.

### The salt response of stomatal density in oil palm

Stomata is hypersensitive to abiotic and hormonal stimulus (Hedrich and Shabala, 2018; Ku et al., 2018). In *Arabidopsis* and other crops, salt stress induces the reduction of stomatal aperture and stomatal density (Huang et al., 2009; Jossier et al., 2010), preventing plants from water loss during osmotic stresses. Interestingly, we found that salt stress increased stomatal density in oil palm (Figure 5). This physiological trait was only found in other ligneous plant such as *Populus alba L* (Abbruzzese et al., 2009). It would be interesting to test the response of stomatal density in other fruit trees. Furthermore, the stomatal density had no difference between each salt stress group, suggesting that low salinity is enough to activate stomatal development with maximum effect. Our data also provided a possible strategy to increase the salt tolerance of oil palm that by salt acclimation with a low salinity before transplanting to higher salinity. Taken together with our data that oil palm could remove excess salt via the stomata (Figure 1), we hypothesized that oil palm balances the stomatal movement and development in response to salt stress. Salt induced stomatal development, allowing salt discharge by transpiration stream via stomata. At the same time, stomatal aperture was reduced to keep the water in the plant. Our research identified the unique salt response of stomatal density in oil palm and introduce an interesting scientific question whether the different responses exist commonly between ligneous and herbaceous plants.

### Elevated EgSPCH expression in oil palm and *Arabidopsis* led to same output of stomatal development but opposite effect on salt tolerance

SPEECHLESS is a master transcription factor which regulates the expression of thousands of genes (Lau et al., 2014). In addition, it is also a key stomatal initiator (Lampard et al., 2008). In our study, EgSPCH was up-regulated in all the three levels of salt stress, therefore its expression and function in *Arabidopsis* was tested. EgSPCH showed similar function with AtSPCH on stomatal development, suggesting that elevated expression of EgSPCH in both oil palm and *Arabidopsis* led to increased stomatal density (Figure 7). However, the increase of stomatal production induced by EgSPCH resulted in weaker salt tolerance (Figure 7). In our salt assay, EgSPCH putatively bound to 41% of DEGs, most of them were critical transcription factors and key regulators in stress tolerance and cell development (Figure 8).. The function of SPEECHLESS in stomatal initiation was verified in other monocot plants (Wu et al., 2019) and the repression of OsSPEECHLESS by salt was found in rice (Kumar et al., 2013), suggesting that the activation of EgSPCH by salt in oil palm is not monocot specific. Our data that salt induce stomatal production in oil palm is in agreement with a previous study in another ligneous plant *Populus alba L* (Abbruzzese et al., 2009). The phenotype that oil palm remove excess salt via stomata is in accordance with that in halophytes (Chen et al., 2019). Therefore, it was hypothesized that ligneous plants or halophytes whose average salt tolerance are better than herbaceous plants, may have evolved a different regulatory network of SPEECHLESS to produce more stomata in response to salt tolerance. It would be valuable to test the stomatal behavior and the SPEECHLESS expression in more ligneous plants and halophytes. The key to solve the functional evolutionary mechanisms between ligneous and herbaceous plants on cell development and abiotic stress tolerance is identification of the upstream regulators of EgSPCH during salt stress.

### The mechanism of salt tolerance in oil palm

Plants respond to environmental factors rapidly at cellular level. Here, we identified a molecular link that connect salt induced signaling pathways to stomatal development. Based on our data, we proposed a working model where salt stress activates stomatal development through activation of the stomatal initiator SPEECHLESS (Figure 9). Transcriptional activation of *EgSPCH* would lead to higher stomatal density, allowing the salt emission via stomata (Figure 9). In addition, the activation of EgSPCH would regulate multiple biological processes via transcriptional control of mass DEGs (Figure 9).

**Figure 9.**
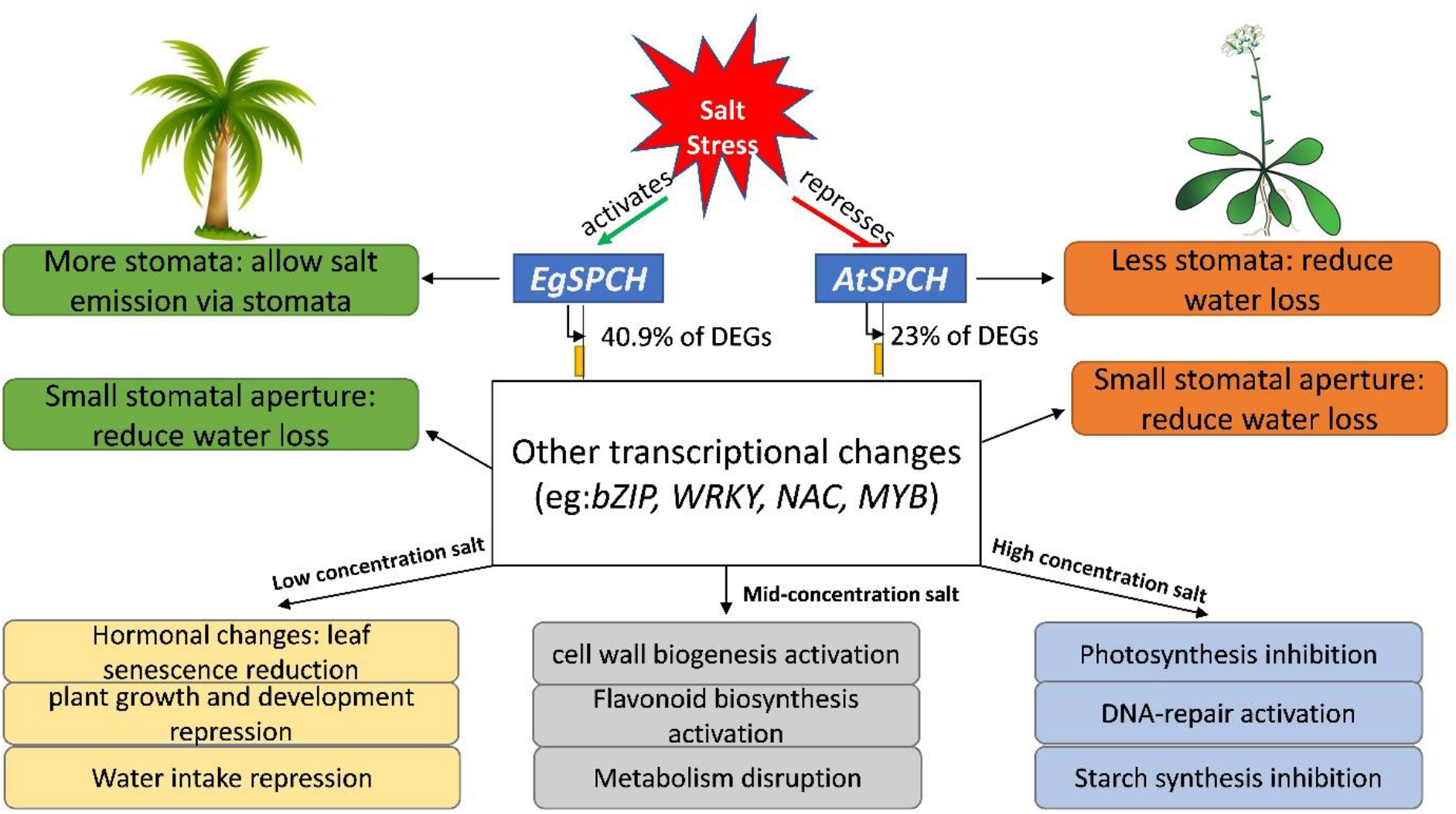
A model of the regulatory networks of oil palm leaves in response to different level of salt stress. In oil palm, salt activates the transcription of EgSPCH in young leaves, which directly increase the stomatal production on leaf epidermis. Furthermore, EgSPCH binds to ~ 41% of DEGs, including key transcription factors that regulate diverse biological processes under different level of salt stress.

Another potential regulation of SPCH expression in oil palm is at the post-translational level via MAPK signaling pathway (Lampard et al., 2008). However, we failed to detect phosphorylated MAPKs using p44/42 MAPK antibody which works well in our previous studies in *Arabidopsis*. Usually SPCH will be regulated with the same direction at both transcriptional and translational level in response to abiotic factors (Lau et al., 2018; Samakovli et al., 2020). The reverse regulation of SPCH at these two levels has not been reported, thus, it appears less likely that SPCH would be repressed in response to salt stress at protein level.

The discovery of salt-induced activation of EgSPCH is novel, as previous studies in rice and *Arabidopsis* either identified the SPCH regulation by MAPKs at the protein level or found the transcription of SPCH is repressed by salt, resulting in less stomata (Kumar et al., 2013; Kumari et al., 2014). Our data explained the phenotype that more stomata is helpful for oil palm to remove excess salt and maintain photosynthesis under salt stress. Nevertheless, the upstream regulatory network of EgSPCH was unknown. Due to the high environmental plasticity of stomata, stomatal assay and transcriptomic analysis in halophytes may be able to answer the question that whether the strong salt tolerance of them are depend on salt-activated SPEECHLESS expression. The comparative genomic analysis and salt stress assay using ligneous and herbaceous crops would be helpful to examine whether the converse salt response of stomata between oil palm and *Arabidopsis* mirrors the different salt tolerance of other crops.

## MATERIALS AND METHODS

### Plant materials and salt treatment

Sixteen two-year-old oil palm seedings with similar sizes were planted in 20 cm diameter pots and were placed in a greenhouse with tropical temperature, 30–50% relative humidity and natural photoperiod. The seedlings were divided into four groups (4 seedlings for each group): Mock group (control group) was watered daily with 150 mL sterilized water while the salt stress groups (100 mM, 250 mM and 500 mM NaCl group) were watered daily with equal volume of 100 mM, 250 mM and 500 mM NaCl diluted by sterilized water, respectively. This is to simulate the condition of the increasing soil salinity caused by mineral weathering or ocean withdrawal. After 14 days of salt stress challenge, the young rosette leaves with similar size in each group were collected.

Col-0 and transgenic *35S:EgSPCH-YFP Arabidopsis* seeds were sterilized and grown on ½ MS plates (0.5 g/L MES, 2.2 g/L Murashige and Skoog salts, 1% [w/v] sucrose, and 0.8% [w/v] agar, pH 5.6) and kept at 4°C in darkness for 3 days. Plants were grown in a well-controlled growth chamber at 22°C with 60% relative humidity under long-day conditions (16 h light/ 8 h dark) at a light intensity of 70 μmol m^−2^ s^−1^. At 7 dpg, 40 well-grown seedlings were transferred to either new ½ MS plates (Control) or ½ MS+ 100 mM NaCl plates.

### Plasmid construction and plant transformation

To generate *35S:EgSPCH-YFP*, the full length CDS sequence of *EgSPCH* was amplified and cloned into pENTR/D-TOPO (Thermo Fisher, USA), after which the entry clone was recombined into the destination vector pGWB541 (Nakagawa et al., 2007) via LR recombination using Gateway LR Clonase II (Thermo Fisher, USA). The primers (EgSPCHcds-F/R) used for plasmid construction are listed in Supplemental Table S1. Transgenic plants were generated in the Col-0 background through *Agrobacterium tumefaciens*-mediated transformation and selected by hygromycin on ½ MS plates.

### RNA extraction and sequencing

Total RNA from oil palm leaves of three biological replicates of control (Mock group) and salt treated samples (100 mM, 250 mM and 500 mM NaCl) was extracted using RNeasy Plant Mini Kit (Qiagen, Germany). RNA quality and quantity assessment, RNA-seq library preparation, library quality control and library quantification were performed using previously described method (Wang et al., 2020). The libraries were sequenced with an Illumina NextSeq500 (Illumina, USA).

### Measurement of stomatal production and stomatal aperture

Small slices from each young rosette leaves collected after 14-day mock or salt treatment were stained in propidium iodide (PI, Molecular Probes, P3566; 0.1 mg/ml) immediately for cell integrity fluorescent microscopy, and images were captured at 20X on a ZEISS Axioscan 7. For quantification of stomatal density and aperture, fresh leaf slices were first cleared in fixing buffer (7:1 ethanol: acetic acid) for 8 hours and were mounted in clearing buffer (8:2:1 chloral hydrate: water: glycerol). Differential contrast interference (DIC) images of the abaxial epidermis of young leaf slices were captured at 20X on a Leica DM2500 microscope. More than 20 slices were examined per test. Stomatal density and stomatal aperture were measured by ImageJ with its built-in tools.

### Differential expressed genes (DEGs) analysis

Adaptor filtering and cleaning of raw sequencing reads were carried out using SeqKit (Shen et al., 2016). Cleaned reads were aligned and mapped to the oil palm reference genome (Singh et al., 2013; Jin et al., 2016) with improved annotation using STAR (Dobin et al., 2013). The expression level of each gene was counted using HTSeq-count (Anders et al., 2015) and the relative expression of each gene was normalized using DESeq2 (Love et al., 2014). Transcripts with more than two times of fold change (FC) value and a significance value less than 0.05 were considered as differentially expressed genes, between mock and salt treatment groups.

### Functional annotation of DEGs

The Gene Ontology (GO) accessions of DEGs were retrieved from the PalmXplore database of oil palm (Sanusi et al., 2018). Principal component analysis (PCA), heatmap analysis, gene ontology enrichment analysis and signaling pathway clustering of candidate genes based on the relative expression of DEGs were performed with the program iDEP (Ge et al., 2018) by referencing to *Arabidopsis*.

### Validation of RNA-Seq data using qPCR

The relative expression of EgSPCH and 11 randomly selected DEGs were tested by qPCR, to examine the validity of the RNA-Seq dataset. The primers used for qPCR are listed in Supplemental Table S1. β-tubulin gene was used as housekeeping gene (internal control) to normalize the relative expression of genes. RT-qPCR was performed in CFX96 Touch Deep Well Real Time PCR System (Bio-Rad, USA) with the program in previous study (Liu et al., 2020). Each gene for qPCR was performed by a biological/experimental triplicate.

## Data availability

Raw RNA-seq reads used in this study have been deposited to the DDBJ DRA database with a DRA submission no. DRA013127

## Supplemental Data

**Supplemental Table S1**. Primers used for plasmid construction and Q-PCR in this study

**Supplemental Table S2**. Selected DEGs and their KEGG pathways in response to 250 mM NaCl challenge in the young rosette leaves of oil palm seedlings

**Supplemental Table S3**. Selected DEGs and their KEGG pathways in response to 500 mM NaCl challenge in the young rosette leaves of oil palm seedlings

**Supplemental Table S4**. All the DEGs and their annotations in each group

**Supplemental Table S5**. Transcription factors among DEGs

**Supplemental Table S6**. Putative EgSPEECHLESS targets among DEGs

**Supplemental Table S7**. Qualities of clean reads from RNA-Seq

**Supplemental Figure S1**. Verification of NaCl crystals on oil palm leaf surface.

**Supplemental Figure S2**. The diversities of differentially expressed genes (DEGs) in response to 250 mM and 500 mM NaCl Supplemental Figure S2.

**Supplemental Figure S3**. Validation of RNA-Seq by Q-PCR.

## Acknowledgements

This work was supported by the Internal Funds of the Temasek Life Sciences Laboratory

---

**Supplemental Table S1.**
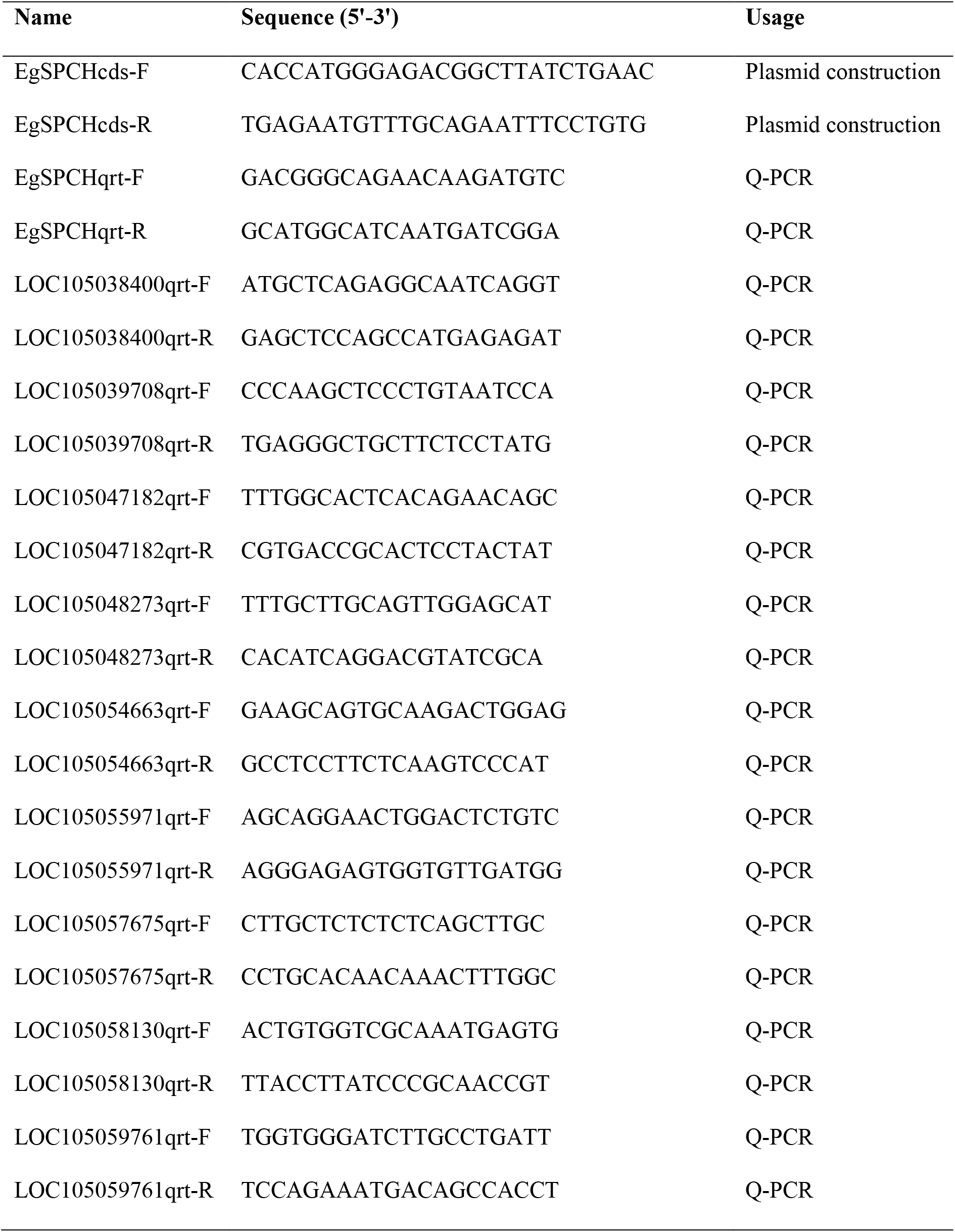
Primers used for plasmid construction and Q-PCR.

**Supplemental Table S2.**
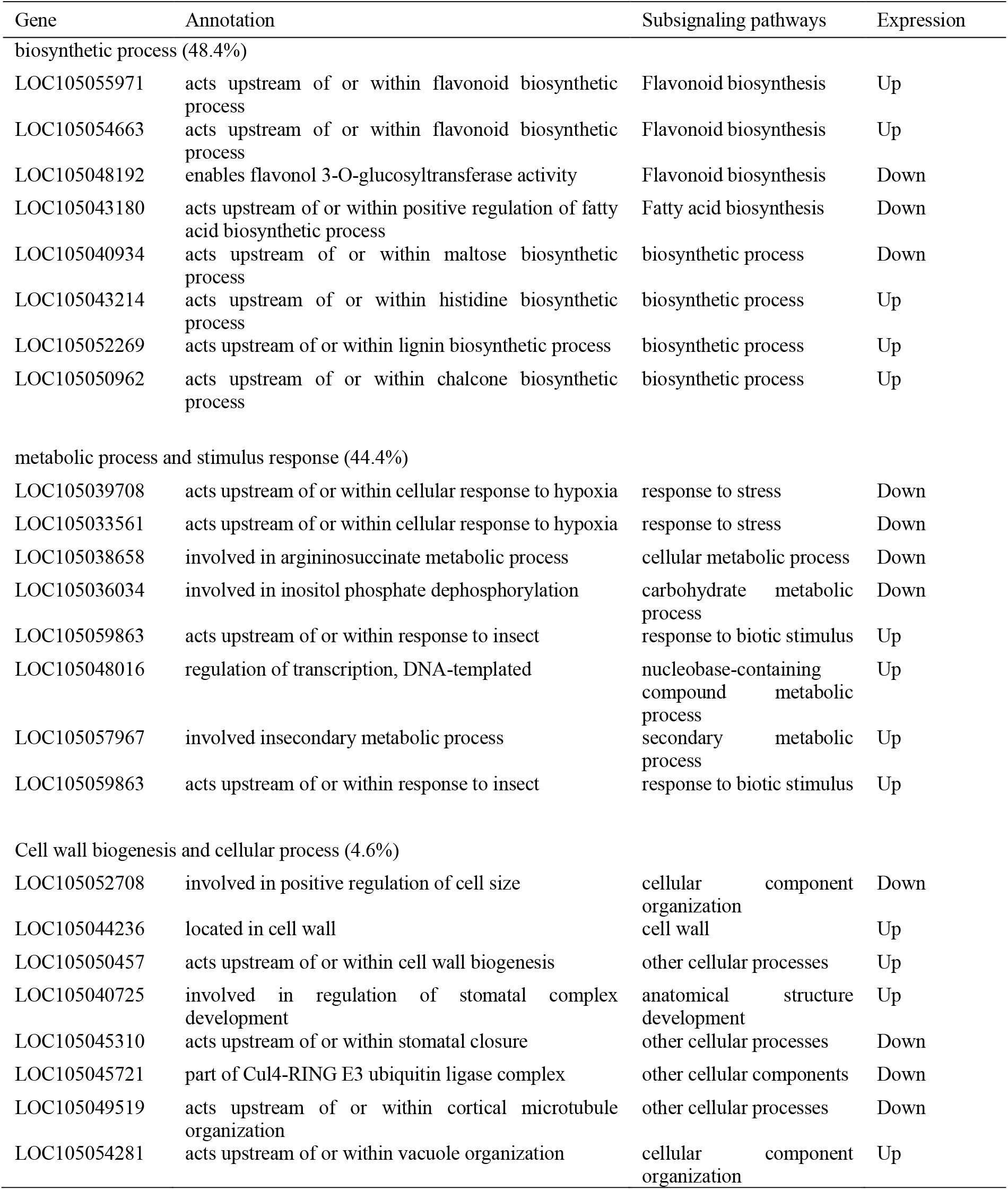
Selected DEGs and their KEGG pathways in response to 250 mM NaCl challenge in the young rosette leaves of oil palm seedlings

**Supplemental Table S3.**
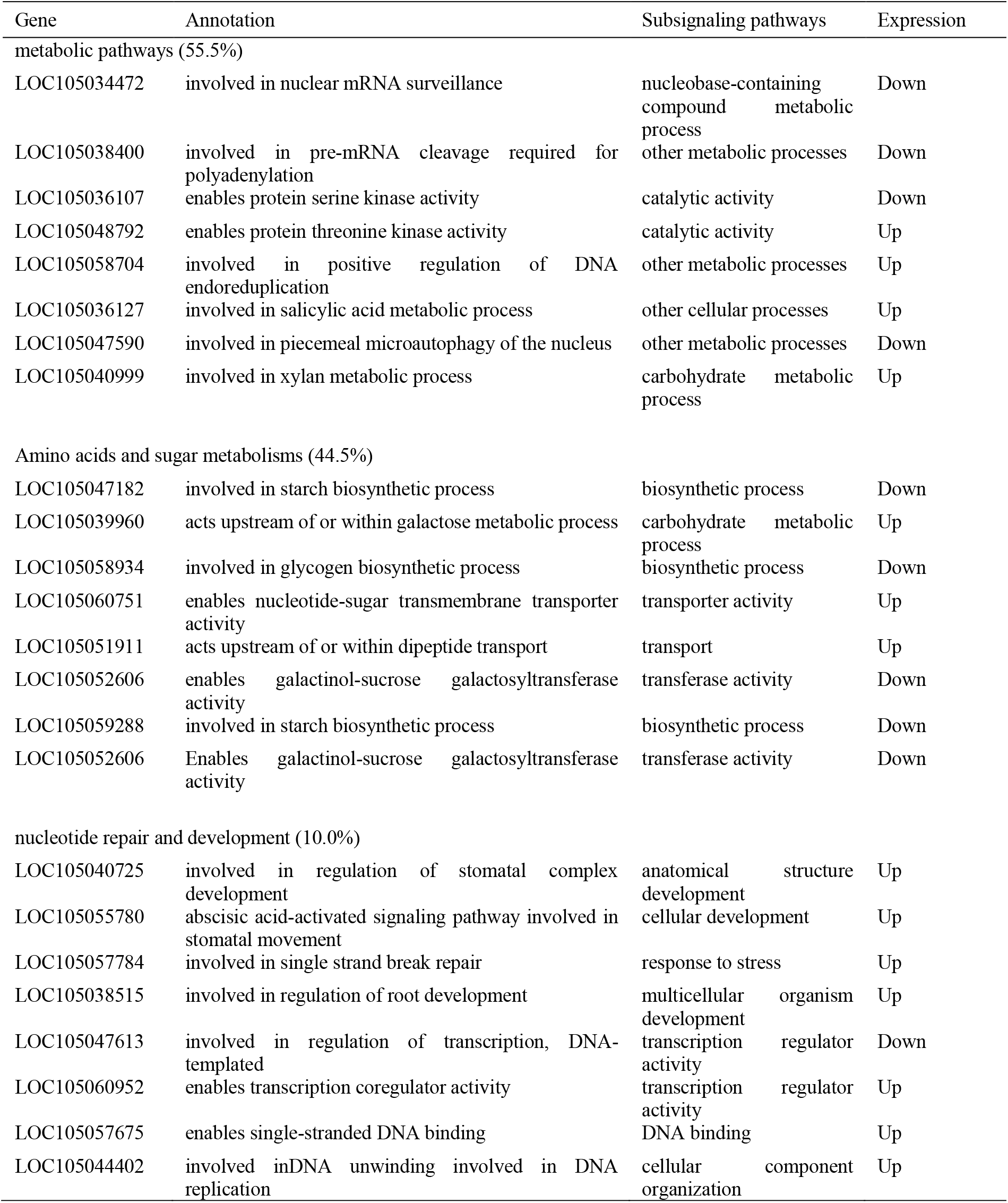
Selected DEGs and their KEGG pathways in response to 500 mM NaCl challenge in the young rosette leaves of oil palm seedlings

**Figure S1.**
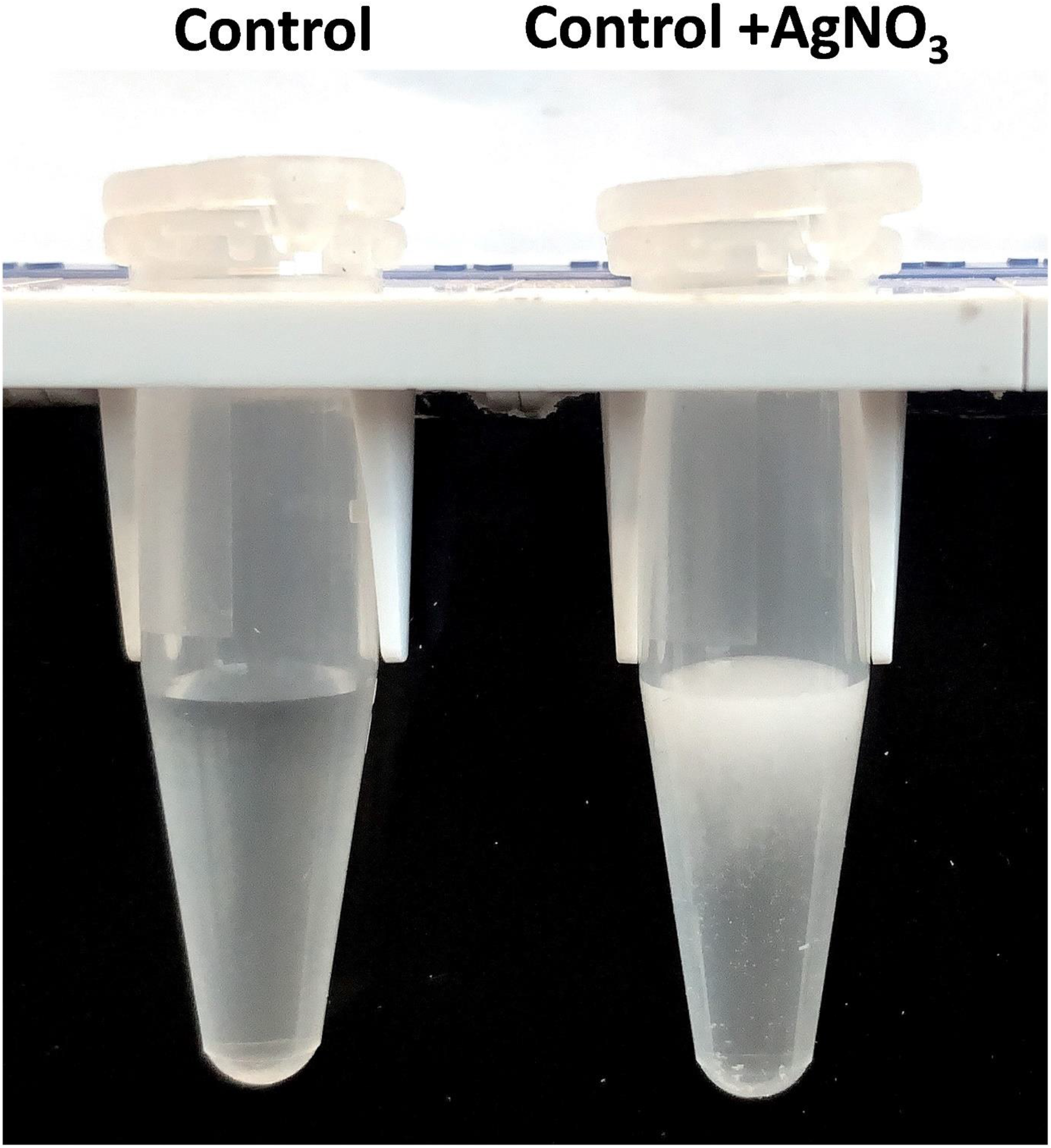
Verification of NaCl crystals on oil palm leaf surface.

**Figure S2.**
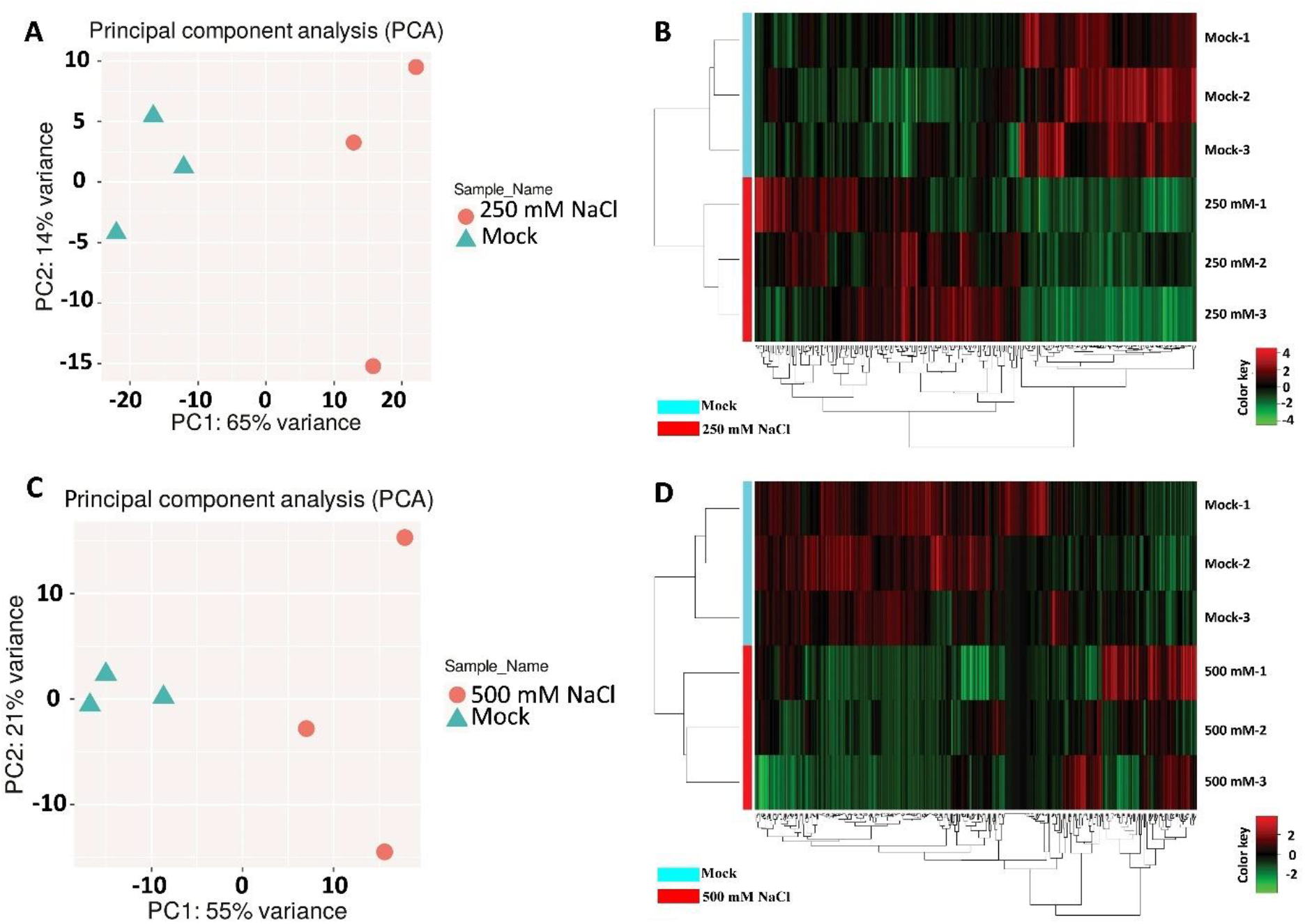
The diversities of differentially expressed genes (DEGs) in response to 250 mM and 500 mM NaCl.

**Figure S3.**
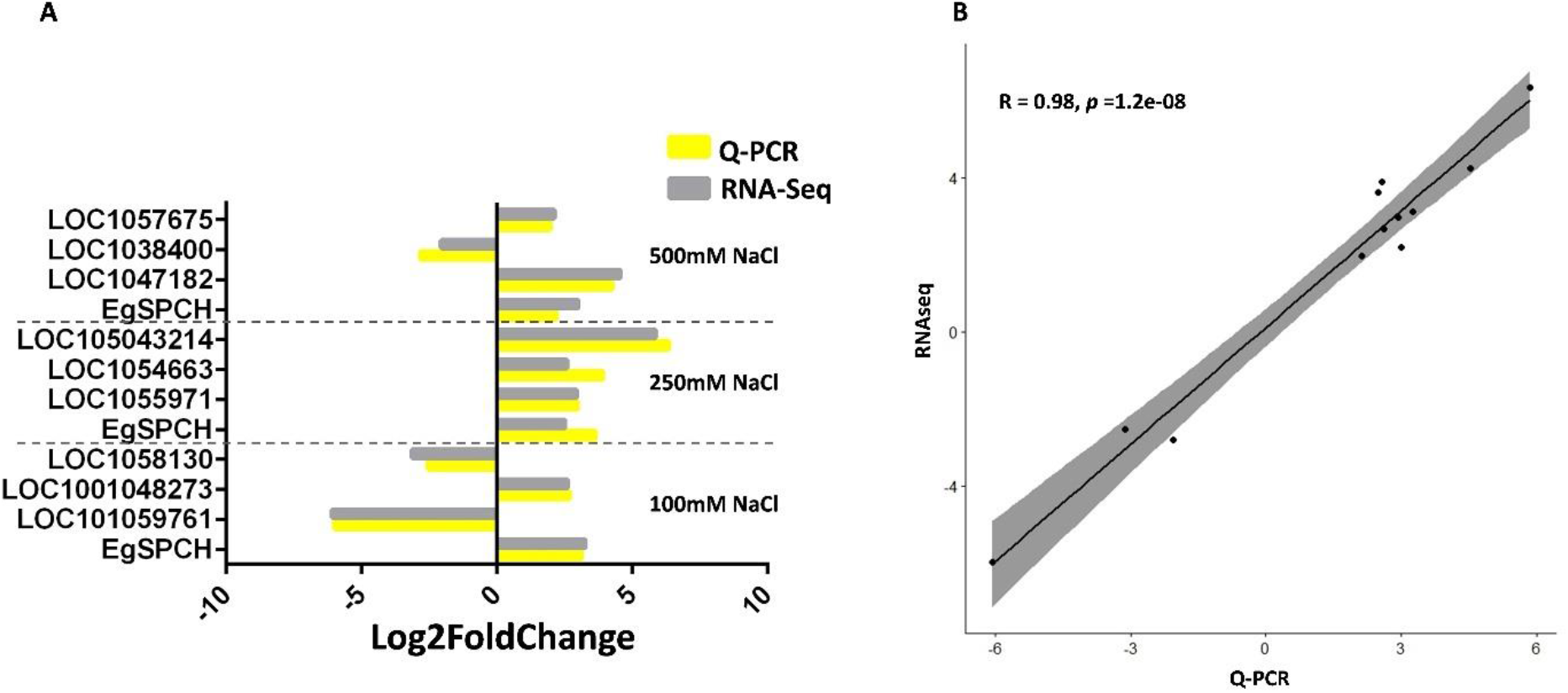
Validation of RNA-Seq by Q-PCR.

